# Behavioral Context Shapes Sensory Responses in Vibrissal Motor Cortex

**DOI:** 10.64898/2026.04.28.720072

**Authors:** Florian Freitag, Jelte de Vries, Liv Grete-Harder, Matthew E. Larkum, Robert N. S. Sachdev

**Affiliations:** Institute of Biology, Humboldt Universität zu Berlin, Charitéplatz 1 / Virchowweg 6, Berlin D-10117, Germany; NeuroCure Cluster of Excellence, Charité–Universitätsmedizin Berlin, Chariteplatz 1 / Virchowweg 6, Berlin D-10117, Germany

## Abstract

Understanding how motor cortical circuits flexibly transform sensory and contextual information into behavior remains a central challenge. Whether neurons in primary vibrissal motor cortex (M1) multiplex across behaviors or are selectively engaged in context-specific actions is still unclear. To address this question, we trained mice on multiple vibrissal sensorimotor tasks, including a cue-triggered whisking-to-touch task and an air-puff–triggered licking task. Fast-spiking and regular-spiking neurons in layers 2/3 and 5 in vM1 responded robustly within ∼15 ms to air-puff stimulation. In contrast, these same neurons were only weakly modulated during goal-directed whisking-to-touch behavior. Unexpected air-puffs evoked responses in fewer neurons than expected stimuli. Trials in which stimulation elicited whisker movements produced smaller neural responses than trials without whisking. Stimulus-evoked activity in M1 was organized along a spectrum of response profiles with neurons exhibiting varying responses dynamics that cut across laminar and physiological distinctions. This organization of responses is consistent with context-dependent recruitment of M1 neurons. Together, these findings indicate that M1 activity is more closely associated with the selection of specific behavioral responses than with generalized sensory-motor encoding.

**Significance:** How motor cortex links sensory input to behavior remains a central question in neuroscience. Do neurons that respond to vibrissal stimuli also participate in whisker-based behaviors, or do they reflect distinct functional states? Here, we show that activity in vM1 is context dependent. Across multiple behaviors, sensory inputs recruit different neuronal populations depending on behavioral context: air-puff stimuli evoke rapid and robust responses, whereas the same neurons are only weakly engaged during goal-directed whisking-to-touch. In addition, expected and unexpected stimuli activate partially distinct ensembles, and sensory responses are attenuated when stimuli directly trigger movement. These findings indicate that M1 does not uniformly encode sensory input; instead, activity reflects context-dependent action selection where neuronal populations are engaged according to behavioral demands.

## Introduction

Neocortical circuits operate in state- and context-dependent regimes that adapt dynamically to behavioral demands (Churchland et al., 2012; Elsayed et al., 2016; Gallego et al., 2017). In motor cortex, neuronal activity reflects movement planning, decision making, and execution, capturing the internal processes through which voluntary actions are prepared and generated (Tanji and Evarts, 1976; Georgopoulos et al., 1986; Fetz, 1992; Li et al., 2015). These actions may be triggered by external sensory events, such as a sudden cue, or arise from internally generated decisions, such as initiating movement in anticipation of an outcome (Evarts and Tanji, 1976; Churchland et al., 2012). Understanding how cortical circuits integrate sensory input with behavioral context to guide action remains a central challenge in systems neuroscience.

The rodent vibrissal system is a model for studying sensorimotor integration (Woolsey and Van der Loos, 1970). Whisker movements actively shape sensory input during exploration (Kleinfeld et al., 2006; Mitchinson et al., 2007; Diamond et al., 2008; Grant et al., 2009). During natural behavior, while navigating and sampling their environment mice whisk at ∼5–20 Hz (Voigts et al., 2008; Sofroniew and Svoboda, 2015; Dominiak et al., 2019; Rodgers et al., 2021). These movements are largely self-initiated and continuous, and whisker–object contact occurs as animals explore their environment. When whiskers contact objects, mechanosensory signals are generated that provide information about object location and shape (Hutson and Masterton, 1986; Petersen, 2007; Diamond et al., 2008; Mitchinson et al., 2011).

Previous studies have shown that whisker stimulation can activate motor cortical areas (Kleinfeld et al., 2002; Ferezou et al., 2007; Huber et al., 2012). Sensory responses in primary motor cortex likely arise through both cortico-cortical projections from primary somatosensory cortex and thalamocortical pathways (Ferezou et al., 2006; Ferezou et al., 2007; Rocco and Brumberg, 2007; Mao et al., 2011; Ohno et al., 2012; Moore et al., 2015; Sreenivasan et al., 2016; Petrof et al., 2015; Pina Novo et al., 2025). Neurons in vibrissal M1 also exhibit activity correlated with whisker movements and tactile stimulation (Carvell et al., 1996; Chakrabarti and Alloway, 2006; Ferezou et al., 2007). Moreover, electrical or optogenetic stimulation of localized regions of M1 can gate sensory processing (Zagha et al., 2013; Urbain and Deschenes, 2007) and drive whisker movements, (Haiss and Schwarz, 2005; Gerdjikov et al., 2013; Friedman et al., 2012; Smith and Alloway, 2013; Chakrabarti et al., 2022).

Despite this extensive literature, it remains unclear whether individual M1 neurons multiplex across distinct whisker-dependent behaviors or whether task context recruits distinct neuronal ensembles. To address this question, we trained mice on two sensorimotor tasks involving whiskers. In one task, an auditory cue paired with an air puff to the whiskers elicited goal-directed licking. In the second task, a different auditory cue prompted mice to actively move a whisker to touch an object. To assess the influence of behavioral context, we also delivered unexpected air-puff stimuli on a small fraction of trials while mice were preparing to whisk. Our recordings targeted a region of mouse M1, that receives input from S1 and corresponds to area RF of vibrissa motor cortex in the rat (Sreenivasan et al., 2016; Haiss & Schwarz, 2005; Gerdjikov et al., 2013), where brief whisker deflections or air-puff stimuli evoke rapid spiking consistent with fast cortico-cortical input from barrel cortex and thalamocortical pathways (Ferezou et al., 2007; Mao et al., 2011; Chakrabarti et al;, 2008; Petrof et al., 2015). Neurons distributed across layers 2/3 and 5 of vM1 responded robustly to expected air-puff stimuli but exhibited weaker responses to unexpected stimuli. Although air puffs rarely evoked whisking, trials in which whisking occurred were associated with reduced stimulus-evoked responses. Across neurons, responses ranged from brief transient increases in firing to sustained activity that outlasted stimulus offset. Critically, most stimulus-responsive neurons did not multiplex across behaviors: neurons activated during the air-puff–triggered licking task were typically not engaged during goal-directed whisking or whisking-to-touch behavior.

## Methods

### Animals and surgery

All procedures were performed in accordance with protocols for the care and use of laboratory animals approved by the Charité–Universitätsmedizin Berlin, Humboldt Universität-Berlin, and the Berlin Landesamt für Gesundheit und Soziales (LaGeSo). Animals were housed in groups of three and kept on a reverse 12-hour day-night cycle to ensure activity during behavioral training and adequate rest.

Adult C57bl6 mice (n=9) mice on a C57bl6 background, weighing 20 to 32 g were anesthetized with Ketamine/Xylazine (90mg/kg / 10mg/kg). Once mice were in the stereotaxic frame (Kopf), a local analgesic Lidocaine (100 μl) was injected under the skin. In the first step, lightweight aluminium headposts were affixed to the skull using Rely X (Applicap, 3M US) followed by Paladur (Henry Schein, UK) (Ebner et al., 2019; Dominiak et al., 2019). Post-surgery analgesia was provided by a combination of Buprenorphine and Carprofen, and in the days after surgery, analgesia was provided by Carprofen (5 mg/kg).

Once animals had been habituated and trained, a second surgery, a craniotomy was made at AP 1.0 and ML 1.0 from bregma. A well was formed by layering Tetric Evoflow A1 (Ivoclar) around the craniotomy on the left hemisphere. Two gold pins were inserted into the well, one as the reference, and one as a backup and secured with additional Tetric Evoflo. A second, smaller craniotomy was performed over the cerebellum and a silver wire soldered to a gold amphenol pin (FHC, Maine) was inserted under the skull. This pin served as the ground for recordings.

### Behavioral tasks

Whisking to touch (W2T): We trained head-fixed mice to use their whiskers to touch a piezo film-based contact sensor. An auditory cue (piezo buzzer) initiated a trial; the cue stayed on for 2 seconds or until the mouse generated whisker contact with the sensor. The contact sensor was placed in front of the right C2 whisker of the mouse, with the animal’s whiskers rostral to the C2 trimmed. The signal from the contact sensor was fed through a custom-built amplifier, the analogue waveform was thresholded to emit a transistor-transistor logic (TTL) signal if contact was made. If whisker-contact evoked a waveform that exceeded a threshold, the auditory cue was turned off and a reward was delivered. The duration of each trial varied and depended on reaction and movement time after cue onset.

Air puff to Lick (A2L): Mice that had been trained in the whisker touch task were also trained to respond to a 250 ms air puff to their whiskers by licking a different “left” lick spout. The air puff was preceded by a piezo buzzer with a different frequency and pattern than the W2T buzzer. The air puff was directed from above the animal’s head in an outward direction, avoiding any direct stimulation of the whisker pad. Mice licked the right lick spout on the W2T trials which were 65% of all the trials and the left spout on the remaining 35% of A2L trials. The skewed distribution of trials was necessary to force mice to attend to and behave for both trials; mice had a preference for the easier trial type where they were expected to respond to air puffs by licking.

Unexpected Air puff (UA): On the experimental day, during recording sessions, on 5 % of trials an unexpected air puff was delivered. The trials began with the auditory cue indicating a W2T trial, but an unexpected air puff (same intensity, location and duration as A2L). Mice were rewarded after the unexpected air puffs.

### Behavioral training

One week after surgery, mice were habituated to being handled and habituated to the behavioral apparatus. In the first days of habituation, mice were acclimated to having their head post handled by the experimenter and were acclimated to several minutes of head fixation. The duration of head fixation was gradually increased from 5 to 40 minutes. During head fixation habituation, mice also became accustomed to having their whiskers puffed, painted, or both. After a week of habituation, mice were water deprived and were trained to respond to an auditory cue and lick the lick tube. The reward was delivered by the opening of solenoid that was audible to mice. Over the course of 2-3 weeks, mice were introduced to the piezo sensor positioned in front of the whisker on the right side of the face. Initially, the sensor was positioned close to the C2 whisker on the right side, with the whiskers rostral to the C2 trimmed back to 2 mm. During the next training days, the sensor was positioned further away from the mouse and the threshold for eliciting reward was set higher -- i.e. mice had to learn to hit the sensor deliberately. To achieve a consistent success rate for each mouse the position of the sensor was moved by a few millimeters from day to day.

### Behavioral tracking

To track whisker and nose positions behavioral data was recorded at 200 Hz with a Basler acA1920-155uc USB 3.0-camera and a f=25mm / F1.4 objective, being set above the animal. The C2 whisker on both sides, as well as the tip of the nose were painted (UV glow, https://www.uvglow.co.uk/). Video data was acquired at 200 Hz. Camera triggers were generated with the reward trigger or at the offset of the auditory cue. A 3.5 second ring buffer was used to record frames of 2.5 seconds before the camera triggers and 1 second after the trigger.

Whisker and nose positions were tracked with Deep Lab Cut from the videos (Mathis et al., 2018; Nath et al.. 2019). For frames where the algorithm failed for any reason (likelihood > 0.9), values were dropped and were filled in later by linear interpolation. Licking was monitored with a piezo sensor attached to the lick tube.

### Electrophysiology

We used two classes of silicone probes to record from Motor cortex in 3 behavioral conditions: Air puff-to-Lick (A2L), Whisking-to-Touch (W2T), surprise Air puff during whisking to touch. The Cambridge Neurotech electrodes (ASSY-77-H5) connected through an Adpt.A64-Om32_2x-sm-NN head-stage to an Intan RHD2164 acquisition board and in separate sessions, Neuropixel 1.0 electrodes connected to a PXIe1000 acquisition card were used to acquire the data. On the day of recording Dura-gel or Kwik cast covering their craniotomy was removed and the area was moistened by filling the recording-well with filtered phosphate buffered saline. A probe dyed with DiI (Thermofisher Vybrant™ Multicolor Cell-Labeling Kit) , a dialkylcarbocyanine (1,1ʹ - dioctadecyl – 3,3,3ʹ,3ʹ - tetra-methylindo-carbocyanine perchlorate), or one of its variants DiO (Snodderly and Gur 1995; DiCarlo et al., 1996; Honig and Hume, 1989) was positioned such that the tip of the probe just contacted the brain. The probe was advanced into the brain slowly at a speed of 5μm per second, to a depth of 900 μm for recordings in cortex; to increase stability Neuropixel 1.0 electrodes were advanced 3400 um into the brain. The probe was allowed to settle for ten minutes. Data was digitized at 30 KHz, and recorded using OpenEphys (Siegle et al., 2017) or SpikeGLX (https://billkarsh.github.io/SpikeGLX/). For Neuropixel 1.0 recordings, the ∼150 channels positioned in cortex were used to acquire data spanning all cortical layers. At the end of each recording session, the probe was slowly (5μm per second) retracted from the brain and the craniotomy was sealed with a two-part silicone gel.

### Track Reconstruction

Mice were deeply anesthetized with isoflurane and transcardially perfused with 0.1% PBS and subsequently with 4% paraformaldehyde (PFA) in 0.1M phosphate buffer (PB, pH 7.2). Brains were carefully removed from the skull and post-fixed in 4% PFA overnight at 4 °C. Brains were embedded in 3% agarose, for sectioning with a vibratome. All brains were sectioned at 70 µm. Before the sections were cover-slipped with ROTI Mount FluorCare mounting medium, sections were wet mounted on a slide and imaged for track reconstruction. Subsequently, sections were cover-slipped for imaging on a CSU-W1 (Nikon) Spinning disc confocal microscope and images were manually aligned to the Allen Brain Atlas.

### Spike Sorting

The data was pre-processed using SpikeInterface (Buccino et al., 2020). We applied a high-pass filter with a minimum frequency of 400 Hz, performed common median referencing, and phase-shifted the signals to account for sampling time differences between the Neuropixels contact sites. Additionally, we detected “bad channels” based on coherence and power spectral density and interpolated between any identified bad channels. Spike sorting was carried out with Kilosort4 using standard parameters (Pachitariu et al., 2024). Putative single units were curated using quality metrics from the Allen software development kit (https://allensdk.readthedocs.io/en/latest/_static/examples/nb/ecephys_quality_metrics.html). The thresholds for amplitude cutoff ratio were set at < 0.1, for inter-spike interval violations the ratio was set at < 0.5, and the presence ratio was set at > 0.9.

Each unit was manually curated in PHY (https://github.com/cortex-lab/phy). This step was used to exclude noisy units and non-neuronal signals. During manual curation, we specifically assessed the number of peaks, the presence of spatial decay, and the presence of a significant dip at zero in the auto-correlogram. All units that had a presence ratio lower than one were examined more closely. In this step, waveform shapes were compared, and cross correlogram ISI violations were assessed to determine whether the unit should be merged with another. If no matches were found, the unit was discarded. This step was necessary to obtain a proper estimate of firing rates in trial averaged plots. In a few recording sessions there were no good units remaining after these quality control steps and the few units that were acceptable had no response to Air puff. We excluded these sessions from further analysis. Taken together, we excluded 3 sessions, 2 with 0 remaining units and one with 5 good units but no responses to stimuli.

### Spike shape

High quality units were classified into three groups, with a manual curation that initiated the process. Regular spiking (likely somatic) had an in initially negative peak while inverted showed an initially positive peak. Units with a trough to peak ratio of less than 0.425 ms calculated using Spikerinterface function (ttps://spikeinterface.readthedocs.io/en/latest/modules/postprocessing.html#template-metrics) were classified as fast spiking, putative interneurons (Senzai & Buzsáki, 2017).

### Statistics

To classify and assess the strength of neural responses, a generalized linear mixed model (GLMM) was used. Z-scores of neural responses were skewed and were therefore modeled using a Gamma distribution. To meet the assumptions of a Gamma GLMM, the data were transformed to include only positive values:

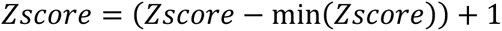

This transformation preserved the shape and relative differences of the distribution. The GLMM was fitted using the glmer function from the ‘lme4’ package in R (Bates et al., 2015). As an example, for Figure 2 E Trial type, cortical layer, and their interaction were included as fixed effects. To account for repeated measures and the nested data structure, random intercepts were included for each mouse and for sessions nested within each mouse. The model used a Gamma distribution with an inverse link function to accommodate the skewed nature of the response variable. Following model fitting, we tested for group differences using the **‘**emmeans**’** package in R, applying multiple comparison correction if necessary (Wickham et al., 2019). All statistical analysis used a similar model structure with the parameters summarized in table 1.

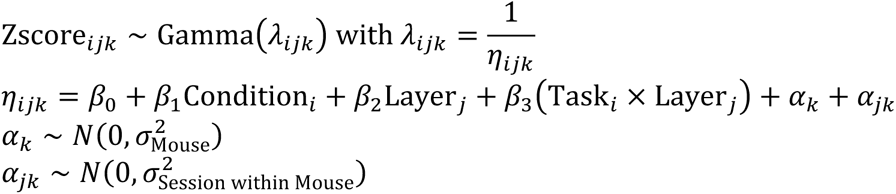

**Table 1.**
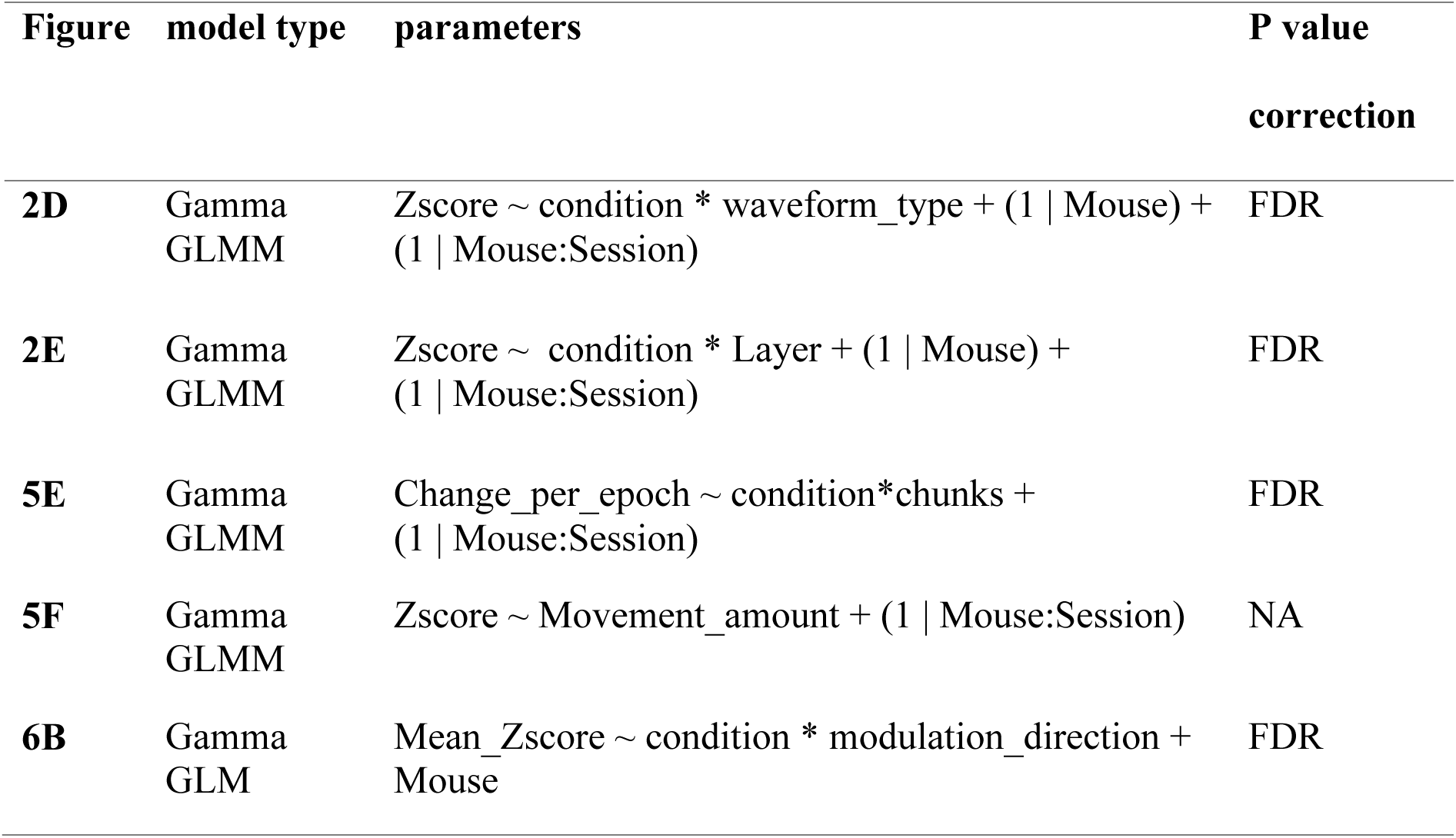
Summary of the model parameters used for statistical analysis. The ‘*’ notations mean the main effects and their interactions. Fixed effects are shown without brackets and random effects are in brackets. Zscore is the max of the Zscore per unit in the first 40 ms post stimulation, Change-per-epoch is the average whisker movement per Behavioral epoch, expressed as percentage of total movement across the trial. Mean_Zscore is the average Zscore across trials and units. The nested random effect was simplified to 1|Mouse:Session when the full model had convergence issues due to insufficient number of samples.

### Significance testing for assessing responsiveness

We assessed neuronal responsiveness using the ZETA-test (Zenith of Event-based Time-locked Anomalies), a parameter-free statistical method designed to detect whether an individual neuron’s firing is significantly time-locked to passive air-puff sensory stimuli or to behavioral events (Montijn et al., 2021). For each neuron, spike times across trials were aligned to the onset of the event of interest, and a cumulative distribution of aligned spike times was constructed. This distribution was compared to a uniform null distribution, and the point of maximal deviation between the two—termed the “zenith”—was identified as the test statistic. Statistical significance was determined by comparing the observed deviation to a null distribution generated by jittering event times across trials, thereby preserving the overall temporal structure of spike trains while disrupting event alignment. For each condition, the analysis was restricted to a 300 ms window relative to behavioral events: 300 ms post-stimulus for Air-puff to lick and Unexpected Air-puff trials, and 300 ms post sensor contact for HIT trials, and 300 ms pre sensor hit for whisking (Whisk). To account for testing multiple units under 4 conditions, we applied the benjamini-hochberg p-value correction. The ZETA-test is available here https://github.com/JorritMontijn/zetapy.

#### Uniform Manifold Approximation Projection (UMAP)

For each unit, spike trains were aligned to air-puff onset and averaged across trials to generate peristimulus time histograms (PSTHs) binned into 10 ms bins. Units responsive to air-puff stimulation were selected for analysis. To normalize for differences in overall firing rate and variance across neurons, firing rates were z-scored separately for each unit within each behavioral condition. PSTHs were smoothed using a rolling mean and normalized within unit to emphasize temporal response shape rather than absolute firing-rate magnitude. Each unit’s normalized PSTH was treated as a series of values across time bins. To reduce dimensionality and denoise the data, principal component analysis (PCA) was applied to the PSTH matrix, and the first five principal components (accounting for 98% of the variance) were retained. K-means clustering (k = 4) was then performed in the reduced PCA space using Euclidean distance to group units according to similarity of their temporal response profiles. For visualization, Uniform Manifold Approximation and Projection (UMAP; n_neighbors = 15, min_dist = 0.1) was applied to the same PCA-reduced data to project high-dimensional activity patterns into two dimensions (McInnes et al. 2018). Cluster membership derived from k-means in PCA space was displayed on the UMAP embedding.

#### PSTH feature extraction

To aid interpretation of the low-dimensional embedding, several scalar features were extracted from each PSTH, including the response integral in early (0–100 ms) and later (100–400 ms) windows following air-puff onset, time to peak response, peak response amplitude, For visualization, feature values were converted to ranks across neurons to reduce the influence of extreme outliers and emphasize relative differences in temporal dynamics. To assess whether anatomical, physiological, or behavioral variables were associated with specific regions of the embedding, neurons were colored by laminar position, spike waveform class, and behavioral modulation (ZETA significance). Distributions of these variables were compared across k-means partitions and across the embedding as a whole.

## Results

### Task design and recordings from primary motor cortex

To determine whether neurons in primary motor cortex (M1) multiplex across sensory-evoked and goal-directed whisker behaviors, we trained mice to perform two distinct tasks that were randomly interleaved within the same session (**Figure 1**). In the *air-puff–to-lick* task, a specific auditory cue was followed by a brief air puff to the whiskers, prompting mice to lick a reward spout. In the *whisking-to-touch* task, a different auditory cue elicited active whisker movements toward a tactile sensor, followed by licking at a separate reward port (**Figure 1A,B**). Whisking-to-touch trials constituted 65% of trials and air-puff–to-lick trials 35%. On recording days, unexpected air-puff stimuli were delivered on a small fraction of whisking-to-touch trials to assess context dependence and responses to surprising sensory input. Facial and whisker movements were recorded from top and side view at 200 Hz and tracked offline using DeepLabCut (**Figure 1C**). Neural activity was recorded with silicon probes targeting vM1 (**Figure 1D**); a subset of penetrations was labeled with fluorescent dyes and registered to the Allen Mouse Common Coordinate Framework (**Figure 1E**). No differences in response parameters or patterns were observed between labeled and unlabeled tracks. Across 29 probe penetrations in 9 wild-type mice, we recorded 4,528 units, of which 1,312 met quality criteria.

**Figure 1.**
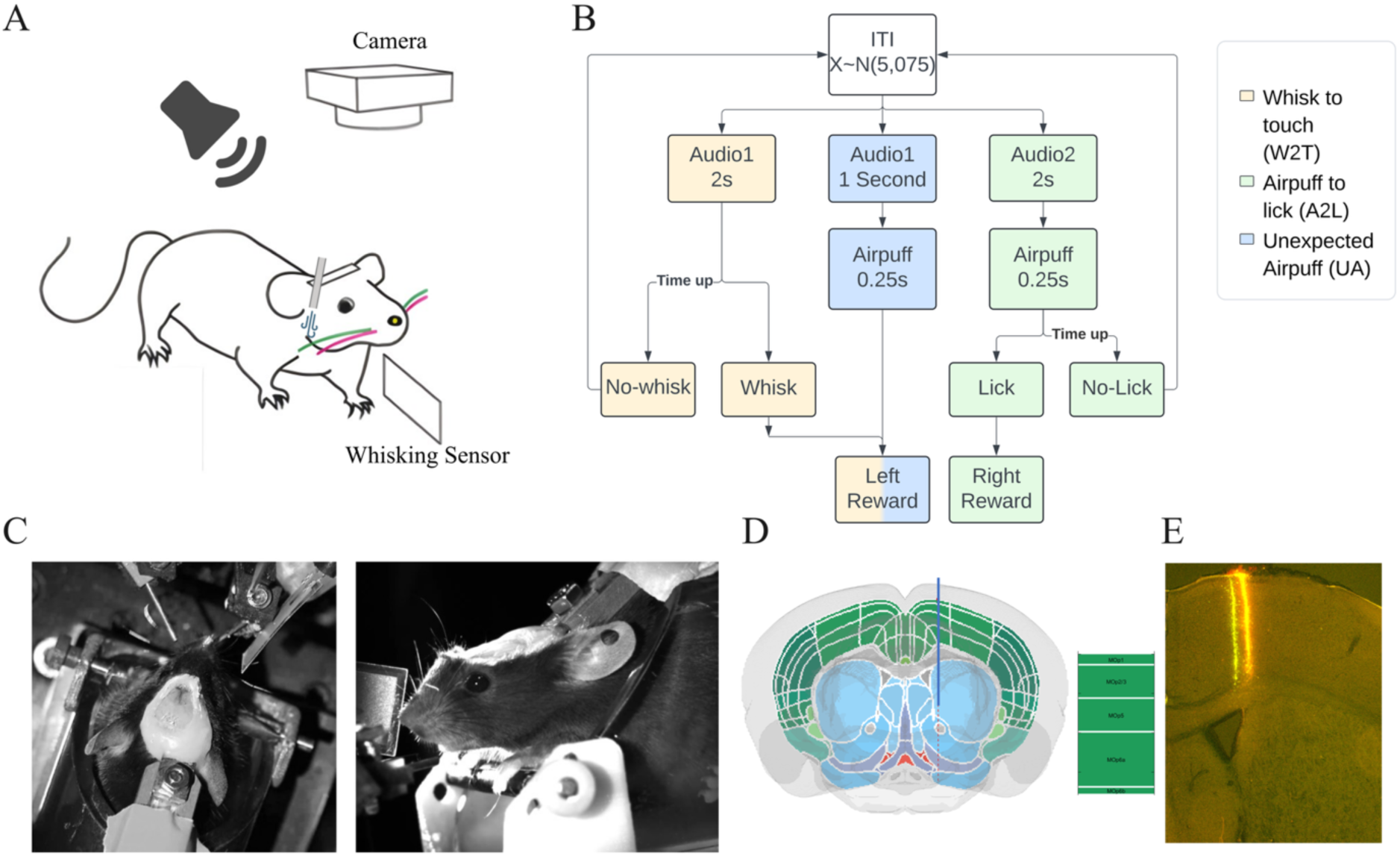
Schematic of experimental approach. **A.** Illustration of head fixed mouse in the whisking-to-touch (W2T) and air-puff to lick (A2L) trials. **B.** Schematic overview of trial types and experimental workflow. Trials were separated by an inter-trial interval (ITI) with a mean of 5 seconds and a standard deviation of 0.75 seconds (*ITI ∼ N* (5, 0.75)). W2T trials occurred more frequently, comprising approximately 60–65% of trials. Each W2T trial began with a 2-second auditory cue, during which mice were required to whisk and make contact with a piezoelectric sensor. Successful contact during the cue period immediately terminated the sound and elicited a reward that was delivered to the left lick spout. Air-puff-to-Lick (A2L) trials were presented less frequently but followed a similar structure. A 2-second auditory cue was paired with a 250 ms air puff. On these trials, mice were required to lick the right spout within 2 seconds to receive a reward. In a subset of W2T trials, an unexpected air puff was delivered early during the trial. In these cases, mice were still rewarded at the left spout following successful sensor contact. **C.** Top and Side view of head fixed mouse. D. Schematic of silicon probe track alignment to the Allen Mouse Brain Atlas in primary motor cortex (M1). E. Representative examples of recovered recording tracks in M1.

**Figure 2.**
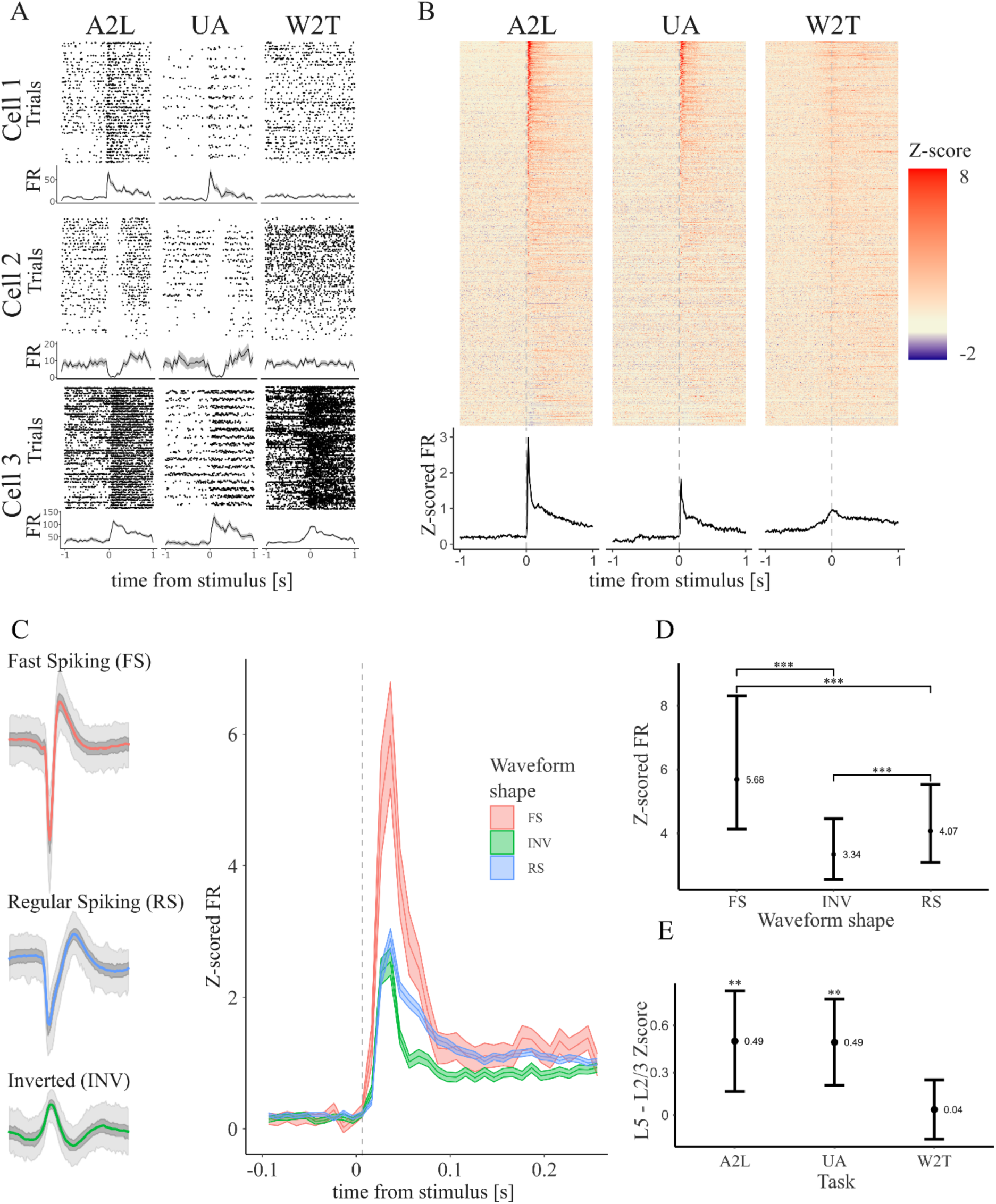
Characterization of M1 responses during active whisking and passive sensory stimulation. **A.** Example rasters and peri-stimulus time histograms (PSTHs) from three single units recorded at different cortical depths, illustrating the diversity of M1 responses across conditions. Responses are shown for three trial types: expected air puffs, unexpected air puffs, and whisking-to-touch trials. The bottom unit displayed increased firing in all three conditions and showed a notable rise in activity prior to time zero, corresponding to whisker movement onset and contact with the sensor. **B.** Population average responses across all recorded neurons for the three conditions: air-puff-to-lick, unexpected air puff during whisking-to-touch trials and whisking-to-touch alone. Air-puff stimuli elicited strong, prolonged responses that outlasted the 250 ms duration of the stimulus. Unexpected air puffs evoked significantly weaker responses compared to expected air puffs. In the whisking-to-touch-task, on average there was an increase in firing before the whisker made contact with the sensor. Shaded region indicates ±SEM. **C, D.** Fast-spiking (FS) neurons exhibited significantly stronger responses to air-puff stimulation compared to regular-spiking (RS) and inverted waveform (INV) units. Data in panel **C** are for Air-puff-to-Lick trials only. Unexpected air-puffs show the same pattern. No statistically significant differences for the different cell types were measured in whisking-to-touch task. Shaded region indicates ±SEM. **E.** Average Response in L5-L2/3. The magnitude of air-puff-evoked responses were slightly but significantly larger for L5 units, compared to responses evoked in L2/3. Units recorded 0-500 microns from the surface were classified as L2/3, and units recorded 500-1200 microns from pia were classified as L5. P-values were calculated from post hoc pairwise contrasts of estimated marginal means based on the fitted generalized linear mixed model and adjusted for multiple comparisons using the FDR method.

### M1 neurons respond robustly to air-puffs but weakly during whisking-to-touch

Example rasters and peri-stimulus time histograms (PSTHs) from single units recorded at different cortical depths illustrate the diversity of M1 responses across task conditions (**Figure 2A**). Air-puff stimuli elicited strong increases in firing in many neurons, whereas activity during whisking-to-touch trials was typically weak or absent. The most common response pattern consisted of robust modulation during both expected and unexpected air-puff stimulation with little or no change in firing during whisking or at whisker contact with the sensor. Less frequent patterns included stimulus-evoked suppression and multiplexed responses in which neurons were active during both sensory stimulation and whisker movement. At the population level, air-puff stimulation evoked a rapid increase in firing beginning approximately 15 ms after stimulus onset, peaking around 25 ms, and decaying over several hundred milliseconds—often outlasting the 250 ms duration of the stimulus (**Figure 2B**). Unexpected air puffs also elicited clear responses, but their magnitude was significantly reduced relative to expected air puffs (p < 0.01, ZETA test). In contrast, activity associated with the whisking-to-touch task increased only modestly prior to sensor contact and remained significantly weaker than responses evoked by air-puff stimulation (p < 0.01). Units were further classified by spike waveform into fast-spiking (FS), regular-spiking (RS), and inverted waveform (INV) units (**Figure 2C**). FS neurons exhibited significantly larger air-puff–evoked responses than RS neurons (24% increase; p < 0.01; **Figure 2D**), consistent with strong recruitment of inhibitory circuitry during sensory stimulation. To determine whether sensory responses varied across cortical depth, units were grouped into superficial (L2/3, 0-500 microns from pia) and deep (L5, 500-1200 microns from pia) layers. Air-puff–evoked responses were slightly but significantly larger in L5 than in L2/3 for both expected and unexpected stimuli (p < 0.05; **Figure 2D**). However, this laminar difference depended on binning parameters and the effect size was modest, suggesting that sensory responses are broadly distributed across layers.

### Context-dependent modulation of M1 responses across cell classes and layers

To examine these results in more detail, and see how behavioral context shaped M1 activity, population responses were compared across the three task conditions—air-puff–to-lick, unexpected air puff (UA), and whisking-to-touch (W2T)—separately for L2/3 and L5 neurons and for FS and RS units (**Figure 3**).

**Figure 3.**
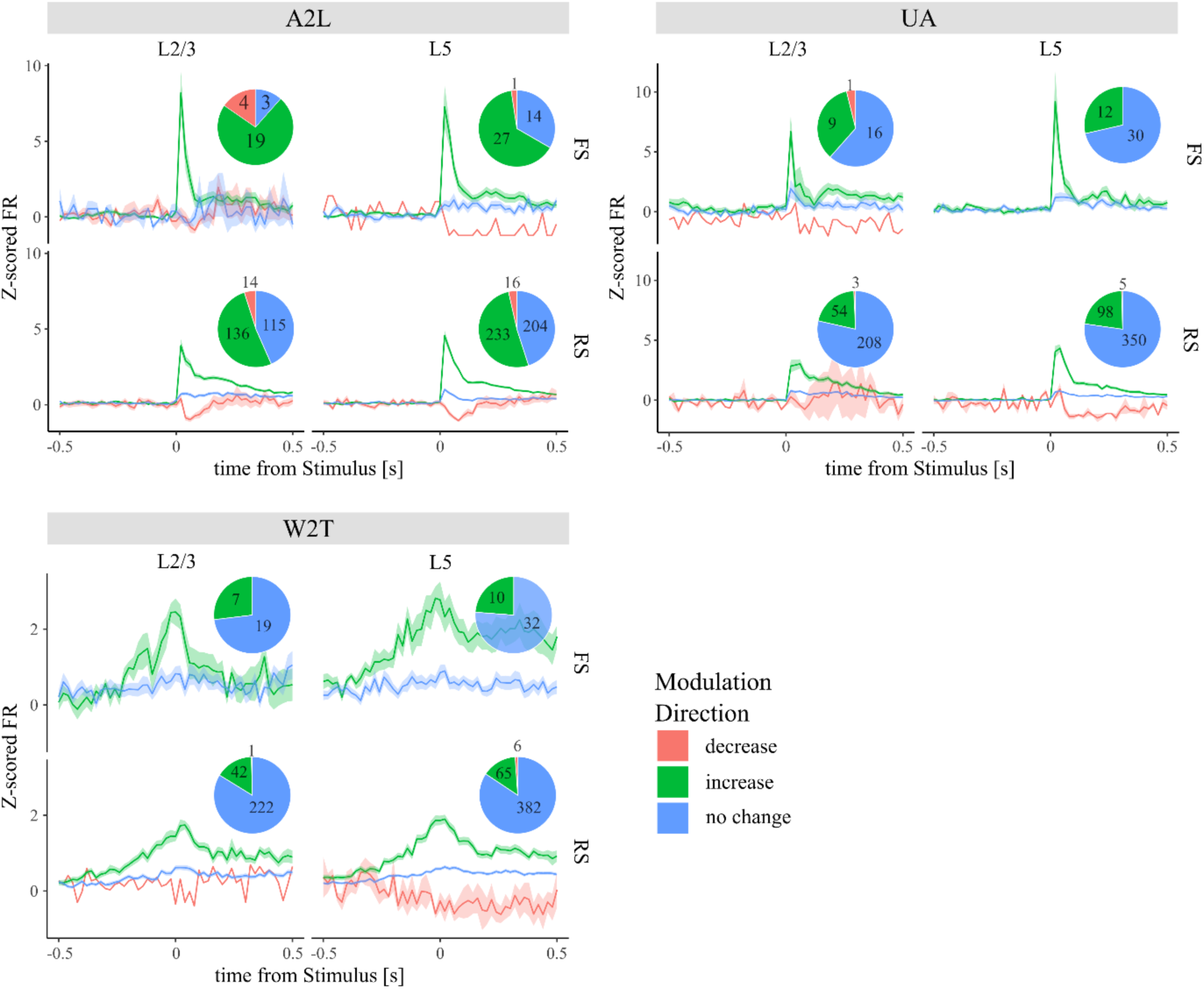
Proportion of distinct response patterns and the average PSTHs of units responding in all 3 behaviors. The z-scored firing rate for 3 patterns of responsiveness: significant increase (green trace), significant decrease (red trace) or no significant change (blue) in response to air puffs for units in L2/3 (left) or L5 (right), for FS (top) or RS units (bottom). Top left panel shows proportion of units responding in Air-puff-to-lick (A2L) trials, Top right shows the proportion of units responding to unexpected air puffs (UA), and the bottom panels shows the response during whisking to touch trials (W2T). Note the massive, 50-60%, reduction in units that respond significantly to air puffs when they are unexpected (compare pie charts in top left and right). Also note relatively weak responses for the same units during whisking to touch trials. Traces show the mean ± SEM across units. Shaded region indicates ±SEM. Pie charts indicate the number of recorded units in each response category and layer.

During the air-puff–to-lick task, both FS and RS neurons showed rapid increases in firing following stimulus onset. Across the population, a large fraction of neurons exhibited significant increases in firing rate, whereas smaller fractions showed suppression or no modulation (**Figure 3**). In contrast, when unexpected air puffs were delivered during preparation for whisking, responses in both layers were reduced and air puffs activated substantially fewer, 50-60% fewer, units. When air puffs were unexpected, or as mice were preparing to move their whiskers to touch, the majority of RS and FS neurons in both layers showed no significant change in firing.

Responses during whisking-to-touch trials differed markedly from those observed during air-puff stimulation. FS and RS neurons in both layers showed weak increases in firing before whisker contact (**Figure 3, bottom**). Across the population, the activity of only 10-20% of all units was significantly modulated during whisking-to-touch, substantially fewer than during air-puff stimulation.

Together, these results show that although many M1 neurons respond strongly to externally delivered whisker stimuli, far fewer neurons are recruited during goal-directed whisking or active touch. Sensory responses in this region of M1 therefore vary strongly with behavioral context and are most robust when stimuli are externally delivered and are expected.

### Dynamics of air puff evoked responses

Air-puff–evoked responses in M1 exhibited substantial heterogeneity in latency, amplitude, and temporal profile. To determine whether this heterogeneity reflected discrete response classes or a continuous organization of response dynamics, we embedded single-unit peri-stimulus time histograms (PSTHs) into a two-dimensional space using UMAP (**Figure 4**). Each point in the embedding represents a single neuron, positioned according to the similarity of its air-puff–aligned firing pattern. The UMAP embedding revealed a curved, continuous manifold rather than well-separated clusters (**Figure 4A**). Applying k-means clustering to the embedding produced partitions that were spatially contiguous but overlapped extensively, without clear boundaries.

**Figure 4.**
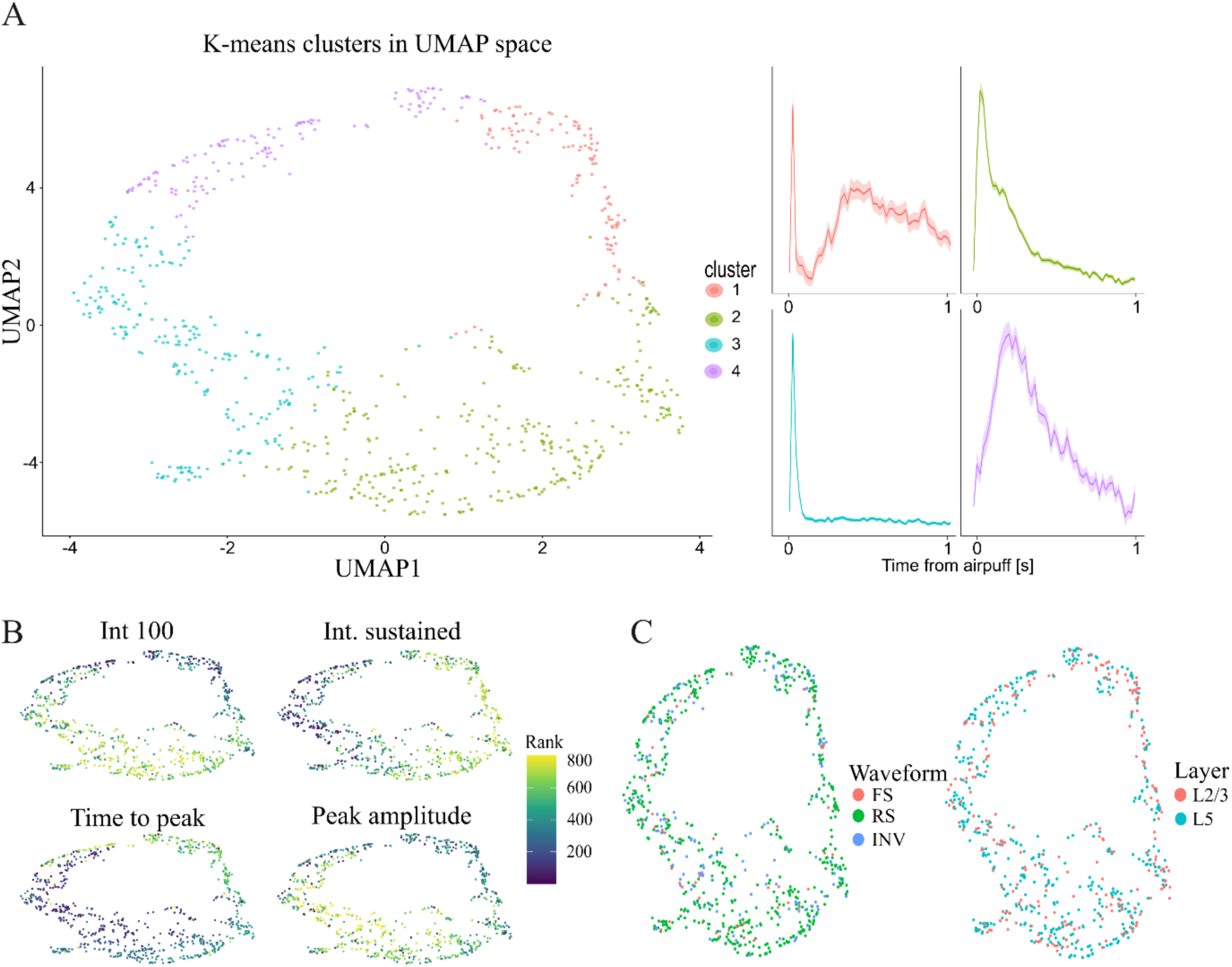
Air-puff–evoked response dynamics in motor cortex. **(A)** UMAP embedding of motor cortex units based on the similarity of air-puff–aligned peristimulus time histograms (PSTHs). Each point represents a single unit. Colors indicate k-means partitions applied to 5-dimensional PCA, showing recurring temporal response motifs. Mean PSTHs for each partition are shown on the right, illustrating characteristic differences in response latency and temporal profile relative to air-puff onset. K-means partitions are shown for visualization and do not imply discrete neuronal classes. Shaded region indicates ±SEM. **(B)** The same UMAP embedding colored by PSTH-derived temporal features, displayed as rank values across neurons to reduce the influence of extreme outliers. Features include the early response integral (0–100 ms), sustained response integral (100–400 ms), time to peak response, and peak response amplitude. Gradual changes in these features across the embedding indicate that the UMAP is organized by temporal aspects of the cue-evoked response, forming a continuous response manifold rather than discrete clusters. **(C)** UMAP embedding colored by spike waveform class (fast-spiking, regular-spiking, inverted) and laminar position (L2/3 vs L5). Neither waveform class nor cortical layer shows systematic alignment with regions of the embedding, indicating that air-puff–evoked response dynamics in motor cortex cut across anatomical and physiological categories.

Examination of the mean PSTHs associated with each partition showed systematic differences in response dynamics, including neurons with sharp, transient responses, neurons with slower and more sustained activity, and neurons with delayed or multiphasic response profiles (**Figure 4A**). However, these motifs transitioned smoothly across the embedding, indicating that they represent graded variations in temporal response structure rather than discrete neuronal categories. To identify the features that structured the embedding, we projected PSTH-derived temporal metrics onto the UMAP space (**Figure 4B**). Early response strength (0–100 ms integral), sustained response magnitude (100–400 ms integral), peak response amplitude, and time to peak all varied gradually across the manifold. Neurons with large, rapid responses occupied regions distinct from those with weaker or more sustained activity, while time-to-peak changed continuously along the manifold. These gradients indicate that the primary axes of organization in the embedding correspond to temporal aspects of the sensory-evoked response rather than categorical differences in response presence or absence.

Next, we examined whether anatomical or physiological properties aligned with specific regions of the response manifold (**Figure 4C**). Coloring the embedding by spike waveform class (fast-spiking, regular-spiking, inverted waveform) revealed extensive intermixing across the entire space, with no class occupying a distinct region. Similarly, neurons recorded in superficial (L2/3) and deep (L5) layers were broadly distributed throughout the embedding, with no strong segregation by laminar position. Thus, neither spike waveform nor cortical depth accounted for the dominant structure of the air-puff response manifold.

Together, these analyses indicate that air-puff–evoked activity in M1 is organized along a continuous spectrum of response dynamics, primarily defined by response latency, amplitude, and persistence. Rather than forming discrete functional classes, M1 neurons exhibit smoothly varying sensory response profiles that cut across laminar and physiological distinctions. This continuous organization is consistent with heterogeneous and context-dependent recruitment of M1 neurons during sensory stimulation, rather than specialized subpopulations dedicated to specific response types.

### Air-puff–evoked whisker movement and its relationship to M1 activity

To determine whether air-puff–evoked activity in motor cortex could be explained by stimulus-induced whisker movement, we quantified whisker kinematics during and after air-puff delivery. In air-puff–to-lick trials, air puffs produced a brief but large, stimulus-locked deflection of the whiskers along both X and Y axes (**Figure 5A**). This movement was transient and largely confined to the stimulus period. Consistent with this observation, plots of the cumulative movement showed that the majority of whisker displacement occurred during the air-puff epoch, with minimal movement during baseline or after stimulus offset (**Figure 5B**). Aligning whisker motion across trials confirmed that frame-to-frame whisker movement increased sharply during air-puff delivery and rapidly returned to baseline immediately after stimulus offset (**Figure 5C**). Thus, in expected air-puff trials, the sensory stimulus reliably deflected the whiskers but did not evoke sustained whisking.

**Figure 5.**
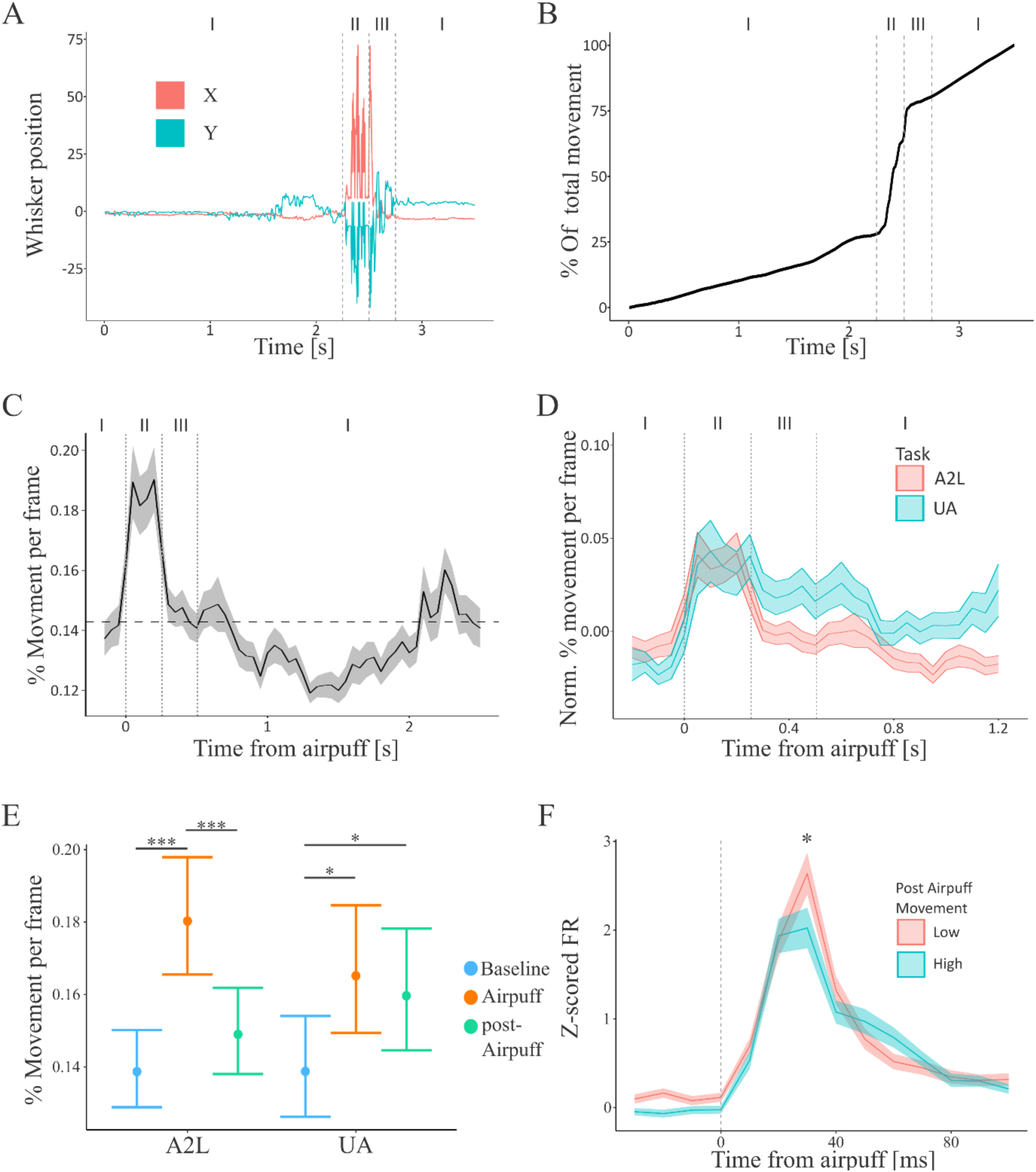
Air-puff–evoked whisker movement and its relationship to M1 activity. **(A)** Example single-trial whisker trajectory showing X (red) and Y (cyan) position over time during an air-puff–to-lick (A2L) trial. Roman numerals indicate three analysis epochs: I, baseline period prior to stimulus and an epoch when mice stop licking and whisking to; II, air-puff delivery; III, post-air-puff period. Whisker displacement is transient and tightly time-locked to stimulus delivery. **(B)** Cumulative whisker movement for the same trial shown in (A), expressed as percentage of total movement across the trial. The majority of whisker displacement occurs during the air-puff epoch (II), with minimal contribution from baseline or post-stimulus periods. **(C)** Average frame-to-frame whisker movement aligned to air-puff onset across trials. Movement increases sharply during stimulus delivery and rapidly returns to baseline levels after stimulus offset, confirming that expected air puffs evoke brief whisker deflections rather than sustained whisking. Shaded region indicates ±SEM. **(D)** Average normalized whisker movement aligned to air-puff onset for expected air-puff–to-lick (A2L, red) and unexpected air puffs delivered during whisking-to-touch trials (UA, cyan). While stimulus-locked movement during the air-puff epoch is comparable across conditions, post-stimulus whisker movement is larger following unexpected air puffs, indicating context-dependent modulation of whisking behavior. **(E)** Quantification of whisker movement across behavioral epochs for A2L and UA trials. Air-puff delivery significantly increased whisker movement relative to baseline in both conditions. In A2L trials, movement returned to baseline during the post-air-puff epoch, whereas in UA trials, post-stimulus movement remained significantly elevated. Points represent the estimated marginal means and 95% confidence Interval (p<0.05 GLMM). **(F)** Average air-puff–evoked M1 firing rate for trials grouped by post-air-puff whisker movement magnitude (higher or lower than the median). low vs high movement). Trials with greater post-stimulus whisker movement exhibited a significantly (p < 0.05) reduced peak M1 response to the air puff, despite similar stimulus timing. This negative relationship indicates that prolonged M1 activity following air-puff stimulation is not driven by whisker movement per se. Shaded regions indicate ±SEM. P-values were calculated from post hoc pairwise contrasts of estimated marginal means based on the fitted generalized linear mixed model and adjusted for multiple comparisons using the FDR method (see methods).

Next, we compared whisker movement evoked by expected air puffs in the A2L task with movement evoked by unexpected air puffs (UA) delivered during whisking-to-touch trials. During the air-puff epoch itself, whisker displacement was comparable across conditions. However, following stimulus offset, whisker movement remained elevated in UA trials relative to A2L trials, indicating that unexpected sensory input triggered additional, context-dependent whisker motion (**Figure 5D**). Quantification across trials confirmed this pattern. In both A2L and UA conditions, whisker movement during the air-puff epoch was significantly greater than during baseline. In A2L trials, post-air-puff movement returned to baseline levels, whereas in UA trials, post-stimulus whisker movement remained significantly elevated (**Figure 5E**). These results indicate that whisker motion following air-puff stimulation depends on behavioral context and expectation.

To assess whether post-stimulus whisker movement contributed to M1 sensory responses, we grouped trials by the magnitude of post-air-puff whisker motion and compared air-puff–evoked firing rates. Trials with greater than the median post-stimulus whisker movement exhibited significantly reduced peak M1 responses compared to trials with minimal post-stimulus movement (**Figure 5F**). This negative relationship indicates that changes in M1 activity following air-puff stimulation cannot be attributed to whisker movement. Also note that even though unexpected air puffs elicit great post stimulus whisker movement, the motor cortical responses to the unexpected air puffs were sparser and weaker than the responses to expected stimuli.

Together, these results show that while air puffs induce large but brief whisker deflections, prolonged M1 responses are not driven by sensory-evoked or self-generated whisker movement. Instead, post-stimulus whisking reflects context-dependent behavioral engagement, whereas air-puff–evoked activity in M1 is dissociable from movement and likely reflects sensory or associative processing rather than motor output.

### Limited overlap of M1 responses across sensory and motor epochs

To examine whether individual M1 neurons multiplex across sensory stimulation, active whisking, and goal-directed touch, we quantified response overlap -- this analysis includes all unit classes: FS, RS and inverted and includes all response classes, i.e. whether they increased or reduced their firing -- across the distinct behavioral conditions: air-puff–to-lick, unexpected air puffs during whisking-to-touch trials, active whisking, and sensor contact (Hit). Neurons were classified as responsive in each epoch using the ZETA test and overlap between conditions was summarized across the population (**Figure 6A**). A large fraction of neurons responded during the A2L condition (319 units), consistent with strong recruitment of M1 by sensory stimulation linked to reward. A substantial subset of these neurons also responded during unexpected air-puff trials (148 units), indicating partial overlap between expected and unexpected sensory stimulation. In contrast, markedly fewer neurons responded only during active whisking (15 units) or at the moment of whisker-contact with the sensor (15 units), and only a small number of neurons were responsive across more than two behavioral conditions: for example, 31 units were responsive to expected and unexpected air puffs *and were responsive* when a whisker hit a sensor. Very few units, only 27, were significantly modulated across all four conditions, indicating that widespread multiplexing across sensory and motor contexts was rare.

**Figure 6.**
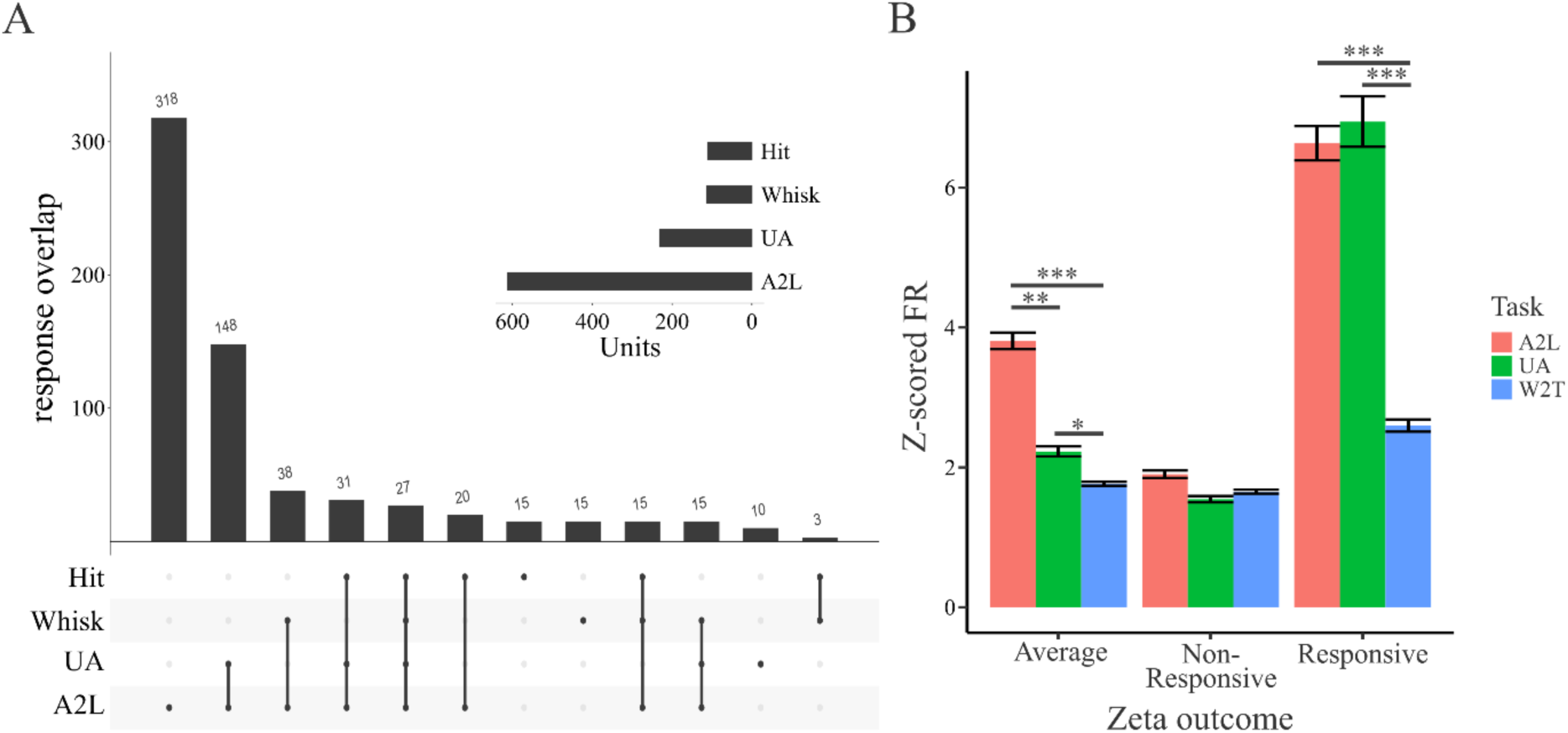
Response profiles of M1 units across behavioral conditions. **A.** Zenith of event based time locked Anomalies (ZETA) test was used to identify M1 units that were significantly responsive during specific behavioral epochs. The inset summarizes the number of responsive units -- including all unit classes -- for each condition: Airpuff-to-Lick, Unexpected Air-puff, Whisking during the Whisking-to-Touch task, and Touch (i.e., sensor contact). The bar graph shows the extent of response overlap, indicating how many units were active in one or more behavioral contexts. Three hundred and eighteen units were significantly active when air puffs were expected in the Air-puff-to-Lick task, while 148 units were active whether the air puffs were during Air-puff-to Lick task or were Unexpected. Thirty-eight units were active during both whisking and in response to expected air puffs. Thirty-one units were responsive when mice whisked to touch and in response to both the expected and unexpected air puffs, and twenty- seven units were responsive during all four behavioral contexts --unexpected air puffs, expected air puffs, whisking and active whisker contact. Most units were selectively engaged by specific behaviors. **B.** Mean z-scored firing rates across behaviors. On average the z-scored firing rate was highest during the Air puff-to-Lick task and lowest during the Whisking-to-Touch condition. Although fewer units responded to the Unexpected Air-puffs than to the expected air puffs, the neurons that were significantly responsive in either condition exhibited comparable Z scores, suggesting similar response strength among recruited units. P-values were calculated from post hoc pairwise contrasts of estimated marginal means based on the fitted generalized linear model and adjusted for multiple comparisons using the FDR method (see methods).

To determine whether differences in recruitment reflected changes in response strength or the number of engaged neurons, we compared average Z-scored peak firing rates across conditions (**Figure 6B**). Across all recorded neurons, A2L trials elicited the largest population response, followed by UA trials, whereas whisking-to-touch (W2T) trials showed the weakest modulation. When the analysis was restricted to neurons classified as significantly responsive within each condition (ZETA-test, with p-values corrected for multiple comparisons using the Benjamini–Hochberg procedure), response magnitudes were comparable between A2L and UA trials. In contrast, neurons responsive during W2T exhibited substantially smaller response amplitudes.

The limited overlap across behavioral epochs demonstrates that M1 neurons that respond to vibrissal stimuli do not multiplex across sensory and motor contexts. Instead, activity in this region of M1 is dominated by condition-specific recruitment, with robust engagement during externally delivered sensory stimuli and sparse engagement during goal directed, self-generated whisker movements and touch.

## Discussion

Motor cortex is often described as integrating sensory signals with motor planning and movement execution. In the vibrissal system, this view predicts that neurons responding to whisker stimulation should also participate in whisker-based behaviors. Our results do not support this expectation. Many neurons in vibrissal M1 responded rapidly and strongly to air-puff stimulation, with firing increasing within ∼15 ms of stimulus onset. However, these same neurons were only weakly modulated during goal-directed whisking-to-touch behavior, and unexpected air puffs evoked weaker and sparser responses than expected stimuli. Thus, neurons that responded robustly to whisker stimulation were rarely engaged during active whisking or tactile contact. At the population level, sensory-evoked and motor-related activity were dissociated, with different behaviors recruiting largely non-overlapping subsets of neurons rather than a shared pool of “whisker-related” cells.

These findings indicate that activity in vibrissal M1 is not organized as a stable representation of whisker stimuli that generalizes across behaviors. Instead, neuronal recruitment depends strongly on behavioral context. This interpretation is consistent with the view that motor cortex operates in context-dependent regimes. Large-scale recordings in primate motor cortex show that the same movement can be represented differently depending on task structure and preparatory state, with population activity reorganizing across conditions (Elsayed et al., 2016; Silvernagel et al., 2026). Our results extend this framework to vibrissal M1: identical sensory inputs engaged partially overlapping but distinct neuronal populations depending on whether stimuli were expected, unexpected or responses were embedded in a goal-directed action where mice whisk to touch a sensor. Thus, rather than encoding sensory events in a behavior-independent manner, M1 appears to operate through context-specific population states that determine which neurons are recruited under different task conditions.

The rapid onset of air-puff responses indicates that sensory signals reach M1 with short latency, likely via fast cortico-cortical and thalamocortical pathways. In our recordings, responses emerged ∼15 ms after stimulus onset and often persisted beyond the 250 ms duration of the stimulus, consistent with previous work in vibrissal sensorimotor circuits (Kleinfeld et al., 2002; Ferezou et al., 2007; Mao et al., 2011; Petrof et al., 2015). This rapid engagement demonstrates that M1 has direct and reliable access to whisker-related sensory input. However, the weak recruitment of these same neurons during goal-directed whisking and the reduced responses to unexpected stimuli indicate that the presence of sensory input alone is not sufficient to drive M1 activity. Instead, these results suggest that sensory signals are selectively gated or routed depending on behavioral context. Under this view, differences in M1 activity across conditions reflect context-dependent modulation of how incoming sensory information is integrated within local circuits, rather than differences in the availability of sensory drive.

Our work aligns with and extends earlier work that is showing how motor cortex engagement changes with behavioral context and the structure of sensory input. Activity in whisker motor cortex (vM1) is strongly driven by input from somatosensory cortex and contributes causally to the initiation of whisking, suggesting a close coupling between sensory drive and motor output (Sreenivasan et al., 2013). However, this coupling does not imply that sensory inputs are uniformly translated into motor cortical activity across behavioral conditions. Rather, our data indicate that identical whisker stimuli can recruit markedly different M1 populations depending on task context and expectation. This view is further supported by work showing that motor cortex is specifically required for responses to unexpected sensory “visual” perturbations but not for the execution of the same movements in unperturbed conditions (Heindorf et al., 2018). Together, these results suggest that M1 is not a general-purpose integrator of sensory and motor signals; M1 is selectively engaged when sensory input is behaviorally relevant in a context-dependent manner—such as when it signals the need to initiate or update an action. In this framework, the reduced and sparser responses we observe to unexpected whisker stimuli, as well as the weak engagement of sensory-responsive neurons during goal-directed whisking, reflect the gating of sensory inputs according to task demands rather than a fixed sensorimotor transformation.

Our results can also be interpreted in the context of recent work showing that VPm thalamic neurons respond robustly to both active touch and passive stimuli, while higher-order thalamic neurons (POm) preferentially respond to passive stimuli but are largely insensitive to active touch (Sumser et al., 2025). This framework provides a useful lens for interpreting our task in which air-puff stimulation constituted a passive stimulus, while whisking-to-touch reflected active sensing. Even though we did not explicitly frame our paradigm in terms of active versus passive touch, our data reveal a similar asymmetry at the level of motor cortex: passive whisker deflections that reliably drive behavior evoke strong and widespread responses in M1, whereas active, goal-directed whisker touch engages these neurons only weakly. Notably, we find that this relationship is not fixed, it is strongly modulated by behavioral context and expectation; unexpected stimuli recruit fewer neurons and weaker responses. Taken together this suggests that distinctions between active and passive sensing are not simply inherited from upstream sensory pathways but are dynamically reinterpreted across thalamocortical circuits. Thus, these neurons in vM1 do not passively reflect the sensory origin of input (active vs passive), but instead they selectively amplify or suppress sensory-driven activity depending on whether the input is behaviorally relevant within the current task context.

Motor preparation and the structure of ongoing behavior likely define the conditions under which sensory inputs are gated in M1. In head-fixed mice, whisker asymmetry reflects preparatory state and behavioral context (Dominiak et al., 2019; Bergmann et al., 2022), indicating that the configuration of the vibrissal system prior to stimulus delivery carries information about the animal’s readiness to act. Reduced responses to unexpected air puffs during preparation for whisking-to-touch are therefore consistent with state-dependent gating of sensory input, in which the current task context suppresses or reshapes incoming sensory drive. In addition, whisker-based behaviors in trained animals are highly structured and learned. Mice performing whisker-to-touch tasks develop stereotyped and individualized orofacial movement patterns (Staab et al., 2025), suggesting that task performance relies on established sensorimotor strategies rather than generic whisking. Under these conditions, sensory inputs that are not aligned with the current behavioral strategy may be attenuated, whereas task-relevant inputs preferentially engage the circuit. Together, these observations support a model in which preparatory state, task structure, and learned behavioral strategies determine how sensory signals are routed through motor cortex, shaping which neuronal populations are recruited in a given context.

The relationship between sensory and motor signals in M1 can be further understood in the context of work from primate motor cortex. In primates, M1 neurons often exhibit strong sensory responses during ongoing movement and participate in rapid feedback loops that support online correction (Evarts, 1973; Pruszynski et al., 2011). In this framework, sensory and motor signals are tightly integrated within the same neuronal populations. Our results point to a different operational regime in vibrissal M1. Although sensory input reaches M1 rapidly and drives robust responses under some conditions, its influence on neuronal activity is strongly contingent on behavioral context. Rather than being consistently embedded within movement-related activity, sensory responses in vibrissal M1 appear to be selectively expressed depending on the current task and preparatory state. This suggests that motor cortex can flexibly shift between modes in which sensory inputs are either integrated with ongoing motor activity or segregated into distinct population configurations. From this perspective, vibrissal M1 may operate in a regime where externally driven sensory events and internally generated motor behaviors recruit partially distinct neural populations, reflecting differences in task structure, behavioral demands, and sensorimotor strategy.

The diversity of sensory responses we observed also provides insight into how M1 neurons are engaged within a single sensorimotor transformation. Dimensionality reduction of air-puff–evoked peri-stimulus time histograms revealed a continuous spectrum of response dynamics rather than discrete response types. Neuronal responses varied smoothly in response latency, peak amplitude, and the duration of post-stimulus activity, forming a structured continuum of temporal response profiles. Neither spike waveform nor laminar position segregated this space, indicating that response diversity does not map onto separable anatomical or physiological classes. Instead, M1 activity during air-puff to lick task is distributed across a continuum in which neurons differ in how they participate in the same stimulus–action mapping. These results argue against fixed functional subtypes and instead support graded, heterogeneous recruitment within a shared population.

Motor cortex is often thought to integrate sensory signals with ongoing movement. Our results suggest a more specific role for vibrissal M1. Neurons in this region respond strongly to whisker stimuli, yet their participation during active whisking and tactile events is weak. Instead of representing whisker movements or touch directly, activity in this part of vibrissal M1 appears to reflect the behavioral context in which sensory signals occur and the actions they predict or trigger. In this view, motor cortex may function less as a general sensorimotor representation and more as a circuit that links behaviorally relevant sensory events to specific motor outputs.

## Acknowledgements

We would like to thank the Charite workshop in particular Jan-Erik Ode, Alexander Schill and Daniel Deblitz for machine and electronics work. We would also like to thank Bryan Hooks (Pittsburgh University), Dieter Jaeger (Emory University), Yangfan Peng (Charite-Universitaetsmedicin, Berlin) for their comments on an earlier version of the manuscript, and Koen Seignette (Humboldt Universitaet, Berlin) for his help with aspects of the analysis, and all other members of the Larkum lab for useful discussions about earlier versions of this manuscript.

## Author Contributions

FF, RS, ML designed research; FF, JV, LGH performed research; FF and LGH analyzed data; RS and FF wrote the first draft of the paper; RS and FF edited the paper; RS and FF wrote the paper.

